# Prey resources are more important than climatic conditions for predicting the distribution of a broad-ranged apex predator

**DOI:** 10.1101/2022.04.04.486974

**Authors:** Luke J. Sutton, David L. Anderson, Miguel Franco, Christopher J.W. McClure, Everton B.P. Miranda, F. Hernán Vargas, José J. de Vargas González, Robert Puschendorf

## Abstract

A current biogeographic paradigm states that climate regulates species distributions at continental scales and that biotic interactions are undetectable at coarse-grain extents. However, recent advances in spatial modelling show that incorporating food resource distributions are important for improving model predictions at large distribution scales. This is particularly relevant to understand the factors limiting distribution of widespread apex predators whose diets are likely to vary across their range. The harpy eagle (*Harpia harpyja*) is a large raptor, whose diet is largely comprised of arboreal mammals, such as sloths and primates, all with broad distributions across Neotropical lowland forest. Here, we used a hierarchical modelling approach to determine the relative importance of abiotic factors and prey resource distribution on harpy eagle range limits. Our hierarchical approach consisted of the following modelling sequence of explanatory variables: (a) abiotic covariates, (b) prey resource distributions predicted by an equivalent modelling for each prey, (c) the combination of (a) and (b), and (d) as in (c) but with prey resources considered as a single prediction equivalent to prey species richness. Incorporating prey distributions improved model predictions but using solely these biotic covariates still resulted in a high performing model. In the Abiotic model, Climatic Moisture Index (CMI) was the most important predictor, contributing 80 % to model prediction. Three-toed sloth (*Bradypus* spp.) was the most important prey resource, contributing 57 % in a combined Abiotic-Biotic model, followed by CMI contributing 29 %. Harpy eagle distribution had moderate to high environmental overlap across all prey distributions in geographic space when measured individually, but overlap was substantially lower in environmental space when prey distributions were combined. With strong reliance on prey distributions across its range, harpy eagle conservation programs must therefore consider its most important food resources as a key element in the protection of this threatened raptor.

## Introduction

Within biogeographic theory, climate is hypothesised to be the main driver of species distributions at continental scales (Wiens 2011; Louthan *et al*. 2015). This is evidenced through the fossil record (Davis & Shaw 2001), and recent observed trends (Walther *et al*. 2002; Parmesan & Yohe 2003). However, the relationship between distribution and climate may be either indirect (Rich & Currie 2018), an oversimplification (Dallas *et al*. 2017), or due to historical biogeography (Heads 2015). Whether biotic resources are more important determinants of species distributions than climatic conditions is still a central issue in ecology and biogeography (Andrewartha & Birch 1954; MacArthur 1972; Wisz *et al*. 2013; Araújo & Rozenfeld 2014; Heads 2015). The current paradigm postulates that biotic resources are most apparent at finer geographical scales (Pearson & Dawson 2003; Peterson *et al*. 2011), but this assertion may not apply across all taxa.

Biotic effects may be lost at continental scales due to the coarse-grain extent, commonly termed the Eltonian Noise Hypothesis (ENH, Soberón & Nakamura 2009). The ENH postulates that because biotic interactions occur at a fine-scale individual level, modelling approaches will fail to recognize them when working at coarse continental scales. Alternatively, biotic resources may correlate closely with abiotic factors, thus the biotic signal is lost in abiotic environmental space (Brewer & Gaston 2003). The effect of biotic resources on species distributions can vary markedly across a given species geographic range (Thompson 2005). Even so, the overriding assumption is that biotic resources require a fine-scale spatial structure to be noticeable (Soberón & Nakamura 2009), because by definition biotic interactions occur at the individual level (Anderson 2016). This assumption is applied to multiple biotic interactions such as the presence or absence of mutualists, competitors, and predators.

The relationship between the range limits of animals, such as butterflies and nectivorous birds being driven by the distribution of their food plants, is well established (Wisz *et al*. 2013; Kass *et al*. 2019). However, it is still unclear if the same processes act on the distribution of large vertebrate apex predators with more diverse diets (Sih 2005; With 2019). It is well-known that apex predators can limit the distribution of their prey species (Holt & Barfield 2009). However, an outstanding question for large vertebrates is whether the distribution of food resources limits the distribution of their main consumers (Sih 2005; Aragón & Sánchez-Fernández 2013; Louthan *et al*. 2015; Schweiger *et al*. 2015). The expectation would be for a high overlap between the abiotic conditions in the predator’s distribution and those of its prey. Consumer (i.e., predator) distribution should be nested within their main resource distributions (Holt 1997), but conversely food resource distributions are not reliant on the distribution of their main consumers.

Food resource distributions can be an important predictor for estimating avian distributions at regional or landscape scales (Aragón & Sánchez-Fernández 2013; de Araújo *et al*. 2014; Aragón *et al*. 2018). However, whether the distribution of food resources can successfully predict the presence of a main consumer across continental extents (2000 – 10,000 km) has not been tested specifically for a terrestrial apex predator. The harpy eagle (*Harpia harpyja*) is a large Neotropical raptor with a continental range across Central and South America from southern Mexico to northern Argentina (Vargas González *et al*. 2006; Sutton *et al*. 2022a). Harpy eagles are distributed across lowland tropical forest (Vargas González & Vargas 2011; Miranda *et al*. 2019; Sutton *et al*. 2021, 2022a), and in seasonal forest enclaves (Silva *et al*. 2013). A recent review summarizing harpy eagle diet across its range established a trend towards a semi-specialized diet (Miranda 2015), mainly comprised of arboreal mammals, including sloths, primates, and tree porcupines. However, birds, reptiles, and terrestrial mammals may also be taken, albeit less frequently (Aguiar-Silva *et al*. 2015; Miranda 2018; Miranda *et al*. 2020).

Species Distribution Models (SDMs) are spatial statistical models that establish the environmental range limits of a given species from environmental conditions and resources at known occurrences (Franklin 2009; Peterson *et al*. 2011). SDMs have seen rapid advances over the past 20 years, yet there are still outstanding conceptual and methodical issues that need addressing to improve predictions (Guisan *et al*. 2017). An important current ecological question is whether including biotic interactions in SDMs can increase their predictive power (Wiens 2011; Wisz *et al*. 2013; Anderson 2017; Dormann *et al*. 2018). Incorporating food resource distributions into abiotic SDMs can improve model predictive performance, leading to more realistic predictions at regional scales (Aragón *et al*. 2018; Atauchi *et al*. 2018; Palacio & Girini 2018). Moreover, including biotic resources in SDMs is especially relevant for species ranging over lower tropical latitudes with more benign abiotic conditions (Louthan *et al*. 2015). Indeed, it has long been hypothesised that species range limits in low-latitude areas are driven more by species interactions than climate (Biotic interactions hypothesis, Dobzhansky 1950; MacArthur 1972). However, given that all taxa need suitable resources and conditions to survive, species distributions must be regulated by both conditions and resources regardless of scale (Godsoe *et al*. 2015). Thus, in this tropical forest predator-prey system, biotic resources and abiotic conditions are expected to exert varying but accountable effects on harpy eagle distribution.

Here, we used a hierarchical modelling approach with four SDMs fitted as functions of abiotic conditions and food resource covariates for the harpy eagle using: **(1)** climatic and topographical covariates (Abiotic model), **(2)** solely food resource distribution covariates (Biotic model), **(3)** including food resource distributions individually, and **(4)** as predicted prey species richness (Abiotic-Biotic models). Further, pair-wise niche overlaps in geographical space were calculated and all distributions were correlated to determine commonality in distribution for all species in this predator-prey system. Lastly, using a novel analytical approach, ordination calculated in environmental space was used to determine the overall extent of niche overlap within this predator-prey system. Specifically, we sought to establish if including food resource distributions were more important for predicting distribution at continental scales and meaningfully improved climate-derived model predictions. Further, we quantified the level of niche overlap between the harpy eagle and its main prey in this lowland tropical forest predator-prey system and predicted areas of highest environmental suitability for the harpy eagle and its main food resources.

## Methods

### Occurrence data

We sourced harpy eagle occurrences from the Global Raptor Impact Network (GRIN, McClure *et al*. 2021) a data information system for all raptor species. For the harpy eagle, GRIN consists of occurrence data from the Global Biodiversity Information Facility (GBIF 2019a), which are mostly eBird records (79 %, Sullivan *et al*. 2009), along with two additional occurrence datasets (Vargas González & Vargas 2011; Miranda *et al*. 2019). Food resource occurrence data were compiled from GBIF (2019b,c,d,e,f,g), following the same data cleaning method, using the five most frequent prey by genus (Miranda 2015): three and two-toed sloth *Bradypus* & *Choloepus* spp. (respectively; 53.2 %), capuchin monkey *Cebus* & *Sapajus* spp (8.0 %), howler monkey *Alouatta* spp. (7.3 %), and tree porcupine *Coendou* spp. (5.3 %). Food resources were combined into their respective genera to: (1) obtain a higher number of occurrence records for each model, and (2) as an appropriate broad scale representation of food resource distribution. Two genera were used for capuchin monkey based on a recent taxonomic assessment, with *Sapajus* (or robust capuchin) found south of the Amazon river and *Cebus* (or gracile capuchin) north of the Amazon river (Lynch Alfaro *et al*. 2011). Combined these five food resource genera comprise 73.8 % by frequency and 75.6% of biomass, representing the majority of food resources taken by the harpy eagle across its range.

We cleaned occurrences by removing duplicate records, those with no geo-referenced location and only those occurrences recorded from 1960 onwards, to temporally match the timeframe of the environmental covariates. For all species occurrences, a 5-km spatial filter was applied between each occurrence point using the ‘geoThin’ function in the R package ENMSDM (Smith 2019). Using a 5-km filter approximately matches the resolution of the environmental raster data (∼4.5-km) and reduces the effect of biased sampling (Kramer-Schadt *et al*. 2013). After data cleaning, a total of 1156 geo-referenced records were compiled for the harpy eagle. Applying the 5-km spatial filter resulted in 715 harpy eagle occurrence records for use in the calibration models. Occurrence records used for the food resource species calibration models are given in Table 1.

**Table 1.** Number of unfiltered and filtered occurrences for food resource genera used in the food resource distribution models (GBIFb,c,d,e,f,g 2019).

| Food resource genus | Primary diet type | Unfiltered | Filtered |
| --- | --- | --- | --- |
| Howler monkey *Alouatta* | Folivore/Frugivore | 1841 | 1045 |
| Capuchin monkey *Cebus* & *Sapajus* | Omnivore | 1160 | 706 |
| Three-toed sloth *Bradypus* | Folivore | 547 | 303 |
| Two-toed sloth *Choloepus* | Folivore | 389 | 237 |
| Tree porcupine *Coendou* | Folivore | 269 | 216 |

### Environmental covariates

We downloaded thirty-seven bioclimatic and topographical abiotic raster layers from the WorldClim (v1.4, Hijmans *et al*. 2005) and ENVIREM (Title & Bemmels 2018) databases at a spatial resolution of 2.5 arc-minutes (∼4.5-km resolution). WorldClim variables (*n =* 19) are generated through interpolation of average monthly weather station climate data from 1960-1990, with ENVIREM variables derived from the WorldClim bioclimatic layers. Raster layers for all species were cropped using a polygon consisting of all known range countries (including Formosa, Jujuy, Misiones and Salta provinces in northern Argentina, and Chiapas, Oaxaca and Tabasco states in southern Mexico). This improves model predictive power by reducing the background area used for testing points used in model evaluation (Barve *et al*. 2011; Radosavljevic & Anderson 2014). Before building each model, all covariates were tested for multicollinearity underlying occurrences using Variance Inflation Factor (VIF, Hair *et al*. 2006). VIF is based on the square of multiple correlation coefficients, regressing a single predictor variable against all other covariates. A stepwise elimination of highly correlated variables was used retaining covariates with a VIF threshold of < 10 (Dormann *et al*. 2013), and Spearman’s Correlation Coefficient of *r_s_* ≤ |0.7| retained for consideration as covariates.

We selected environmental covariates for the food resource distribution models based on species biology and reducing collinearity between environmental covariates underlying the occurrences of each specific food resource genus (Meineri *et al*. 2012). Using this method all five food resource models used a different set of environmental covariates (Table S1), resulting in low collinearity between the final food resource model raster covariates (all tests VIF = < 10; Table S2). Collinearity tests were calculated using the ‘corSelect’ function in the R package FUZZYSIM (Barbosa 2015, 2018) and the ‘vifstep’ function in the R package USDM (Naimi *et al*. 2014). For the Abiotic-Biotic SDMs, these five covariates defining modelled food resource distributions were included in the harpy eagle model calibration as raster layers as has been done in previous studies (Arajúo & Luoto 2007; Preston *et al*. 2008; Ghergel *et al*. 2018). Finally, we combined all individual food resource models into a stacked SDM (s-SDM) and the continuous suitability values summed for a continuous estimate of food resource species richness in the range 0.0 – 5.0.

For the Abiotic model (**A**) two climatic variables, Climatic Moisture Index (CMI) and minimum temperature warmest month, and one topographic variable, Terrain Roughness Index (TRI), were included as covariates. We selected all three covariates *a priori* because combined they contributed 96 % to model prediction from a previous SDM (Sutton *et al*. 2021). Food resource distributions were used in three further models, with SDMs built and fitted using the same methodology as for the Abiotic model. First, only food resource distribution covariates were used in a Biotic model (**B**). Second, modelled food resource raster predictions were included as individual biotic covariates along with the abiotic covariates in an Abiotic + Biotic model (**A+B**). Finally, the predicted species richness was used as the sole biotic predictor along with the abiotic covariates in an Abiotic + (Biotic) Species Richness model (**A+SR**), for a comparison to using individual prey genera (**B**) as covariates. Geospatial analysis and modelling was performed in R (v3.5.1; R Core Team, 2018) using the DISMO (Hijmans *et al*. 2017), RASTER (Hijmans 2017), RGDAL (Bivand *et al*. 2019), RGEOS (Bivand & Rundle 2019) and SP (Bivand *et al*. 2013) packages.

### Species Distribution Models

We fitted SDMs using penalized elastic-net logistic regression (Fithian & Hastie 2013), using a point process modelling (PPM) framework in the R package MAXNET (Philips *et al*. 2017). Elastic net logistic regression imposes a penalty (regularization) to the model shrinking the coefficients of covariates s that contribute the least towards zero (or exactly to zero). The complementary log-log (cloglog) transform was selected as a continuous index of environmental suitability, with 0 = low suitability and 1 = high suitability. The MAXNET package is based on the maximum entropy algorithm MAXENT, equivalent to an inhomogeneous Poisson process (IPP; Fithian & Hastie 2013; Renner & Warton 2013; Renner *et al*. 2015). Philips *et al*. (2017) demonstrated the cloglog transform is equivalent to an IPP and can be interpreted as a measure of relative occurrence probability proportional to a species potential abundance.

We used a random sample of 10,000 background points as pseudo-absences recommended for regression-based modelling (Barbet-Massin *et al*. 2012) and to sufficiently sample the background calibration environment (Guevara *et al*. 2018). Optimal-model selection was based on Akaike’s Information Criterion (Akaike 1974) corrected for small sample sizes (AIC_c_; Hurvich & Tsai 1989), to determine the most parsimonious model from two key MAXENT parameters: regularization beta multiplier (β; level of coefficient penalty) and feature classes (response functions, Warren & Seifert 2011). For all SDMs, eighteen candidate models of varying complexity were built by conducting a grid search comparing a range of regularization multipliers from 1 to 5 in 0.5 increments, and two feature classes (response functions: Linear, Quadratic) in all possible combinations using the ‘checkerboard2’ method of cross-validation (*k*-folds *=* 5) within the ENMEVAL package in R (Muscarella *et al*. 2014).

We only used Linear and Quadratic features to produce less complex and more realistic predictions (Merow *et al*. 2013; Guevara *et al*. 2018). The checkerboard cross-validation method of partitioning masks the geographical structure of the data according to latitude and longitude lines, dividing all occurrences into four spatially independent bins of equal numbers. By masking the geographical structure of test-data the models are projected onto an evaluation region not included in the calibration process. All occurrence and background test points are assigned to their respective bins dependent on location, thus further reducing spatial autocorrelation between testing and training localities (Radosavljevic & Anderson 2014). We used response curves, parameter estimates and percent contribution to model prediction to measure variable performance within the optimal calibration models.

### Model evaluation

We evaluated model performance using both threshold-independent and threshold-dependent measures (Radosavljevic & Anderson 2014). Omission rates are a threshold-dependent measure that report the proportion of training points that are outside of the model when converted into a binary prediction. Omission rates evaluate discriminatory ability and over-fitting at specified thresholds. Lower omission rates show improved discrimination between suitable and unsuitable habitats (indicating higher performance), whilst overfitted models show higher omission rates than expected by theory (Radosavljevic & Anderson 2014). A single threshold-dependent measure was calculated based on the 10% training presence omission rate (OR10) threshold. For low over-fit models the expectation for OR10 is a value close to 0.10 (Muscarella *et al*. 2014).

We used Continuous Boyce index (CBI, Hirzel *et al*. 2006) as a threshold-independent measure of how predictions differ from a random distribution of observed presences (Boyce *et al*. 2002). CBI is consistent with a Spearman correlation (*r_s_*) with CBI values ranging from −1 to +1, with positive values indicating predictions consistent with observed presences, values close to zero no different than a random model, and negative values indicating areas with frequent presences having low predicted environmental suitability. CBI was calculated using five-fold cross-validation on 20 % test data with a moving window for threshold-independence and 101 defined bins in the R package ENMSDM (Smith 2019). We evaluated models against random expectations using partial Receiver Operating Characteristic ratios (pROC), which estimate model performance by giving precedence to omission errors over commission errors (Peterson *et al*. 2008). Partial ROC ratios range from 0 – 2 with 1 indicating a random model. Function parameters were set with a 10 % omission error rate, and 1000 bootstrap replicates on 50 % test data to determine significant (α = 0.05) pROC values >1.0 in the R package ENMGADGETS (Barve & Barve, 2013).

### Geographical overlap and correlation

We measured pair-wise geographic overlaps between the harpy eagle and the five prey distributions using Schoener’s *D* (Schoener 1968, Warren *et al*. 2008), which ranges from 0 (no overlap) to 1 (identical overlap). To estimate the areas where all six species coincide, the three harpy eagle SDM predictions that used biotic covariates (B, A+B and A+SR) were first stacked and their respective CBI scores used to calculate a weighted mean ensemble prediction. Second, the five prey distribution SDMs were also stacked into a single raster. Lastly, we then predicted a measure of commonality in species distribution by intersecting the harpy eagle ensemble prediction, with the stacked prey distribution rasters with a threshold of 0.5 using the ‘stability’ function in the R package SDSTAF (Atauchi 2018).

### Environmental overlap

We quantified overlap in environmental space between the harpy eagle and its five main prey species, using a principal component analysis (PCA) to directly compare species-environment relationships in environmental space in contrast to geographical space using SDMs. Using the PCA-env framework of Broennimann *et al*. (2011), we performed an ordination using the R package HUMBOLDT (Brown & Carnaval 2019). We used the same filtered occurrences for the harpy eagle, but all filtered prey species occurrences were combined into a single occurrence dataset. Nine covariates (eight climatic and one topographical) were used from the WorldClim and ENVIREM databases (Table S3). Collinearity was reduced on the entire raster area by removing non-relevant covariates and final predictor selection checked for correlations using the ‘vifstep’ function in the R package USDM (Naimi *et al*. 2014; VIF = < 7).

We set the PCA-env method to calibrate on a non-analogous environmental space using a minimum convex polygon around all spatially filtered occurrences on a 100 × 100 resolution grid, with a smoothed Gaussian kernel density function (bandwidth = 1). Environmental overlap was quantified on the first two principal components using Schoener’s *D* statistic. Using smoothed densities allows measured overlap to be independent of grid resolution, important for unbiased estimates of niche overlap using Schoener’s *D* (Broennimann *et al*. 2011). We tested equivalency in shared environmental space, first using a niche Equivalency Statistic to test for the difference (α = 0.05) between the observed overlap scores and those under a null distribution hypothesis that the two distributions are equivalent (Warren *et al*. 2008). For the null distribution, presence points are randomly assigned to each species, and an SDM is built on these randomized data. This is repeated a hundred times and a probability distribution is then estimated for niche overlap under the null hypothesis that both sets of species’ occurrences are randomly distributed in the environment. Second, to measure the ability of the Equivalence Statistic to detect differences in environmental space, we used a Background Statistic to test for the difference (α = 0.05) if the observed occurrences of one species are more similar than expected by chance to the background occurrences of the other species (*n =* 100; Warren *et al*. 2008).

The background test corrects for the environmental heterogeneity inherent in environmental data underlying occurrence data, assuming that all species are choosing environments at random throughout their respective geographic ranges. If distributions are not equivalent, a statistically significant difference allows rejection of the null hypothesis of niche equivalency between the two distributions, regardless of the significance of the Background Statistic. A non-significant Equivalence Statistic and significant Background Statistic supports the hypothesis of equivalent shared environmental space. If both statistics are non-significant this implies niche equivalency could be the result of shared identical environmental space, with limited power for the Equivalency Statistic to detect any significant differences. Importantly, the Background Statistic assesses the power of the equivalency test by asking if two distribution models are equivalent based on the matching environments available. It does not provide any evidence that niches are not equivalent.

## Results

### Food resource distribution models

Optimal model selection (ΔAIC_c_ = 0.0) for all prey distribution models had feature classes Linear and Quadratic and a regularization multiplier β = 1, except for two-toed sloth with β = 1.5. Discrimination ability (OR10) for all models was at or close to expected thresholds (Table 2). Final best-fit models were robust to random expectations (range: pROC = 1.344-1.802) with high model calibration accuracy (range: CBI = 0.840-0.969). Capuchin monkey had the broadest distribution, followed by howler monkey and tree porcupine (Fig. 1). Three-toed and two-toed sloths were largely restricted to Central America, Colombia, Amazonia and the Guiana Shield. Prey species richness was highest in Panama, north along the Caribbean coast of Central America, and south along the Pacific coast of Colombia. A broad belt of high prey species richness was predicted across northern Amazonia, east into the Guiana Shield and across the central Amazon.

**Figure 1.**
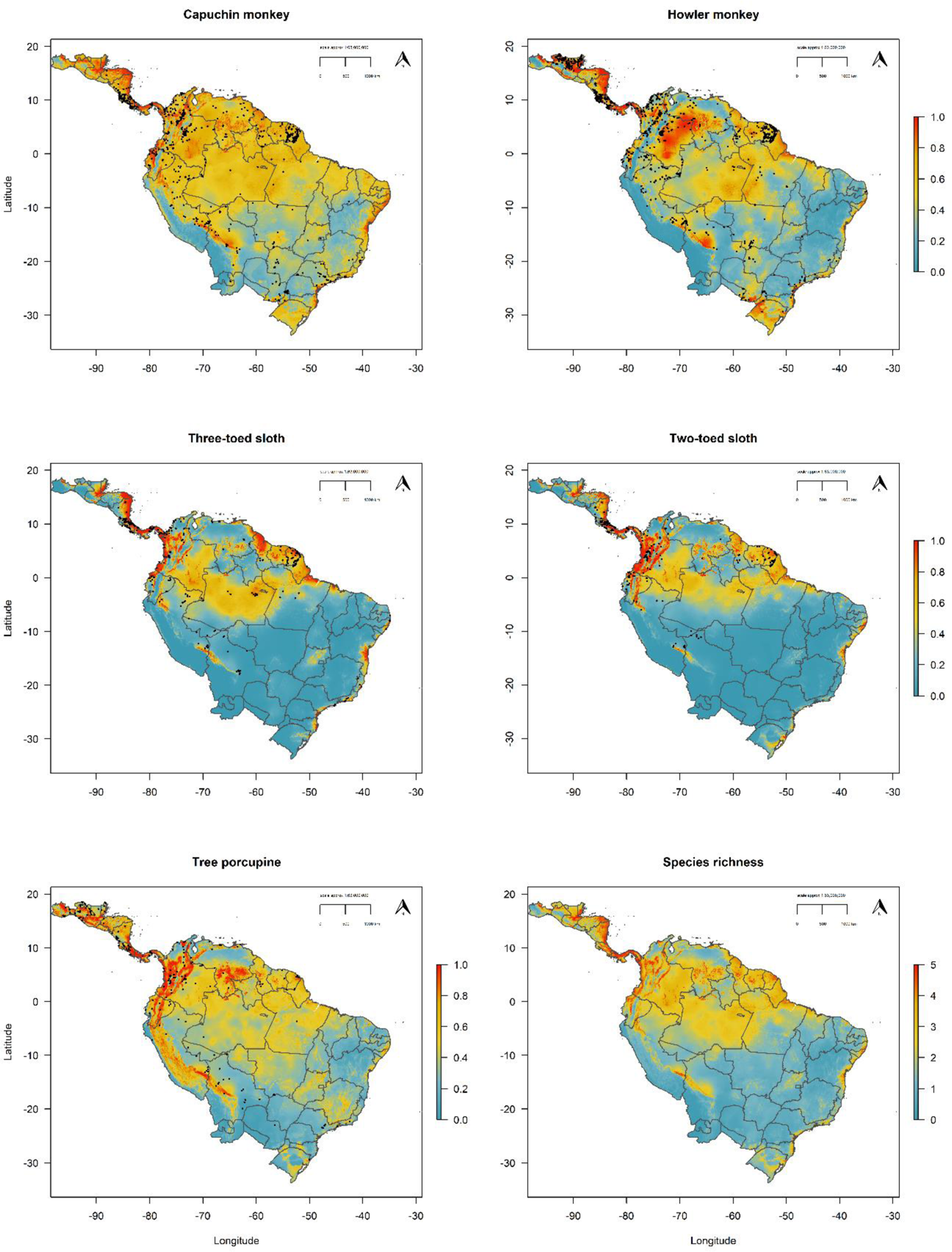
Predicted distributions for the five primary prey genera for the harpy eagle and combined into a summed prediction of prey species richness. Maps denote cloglog prediction with red areas (values closer to 1) having highest suitability. Grey borders represent national borders and state boundaries for Argentina, Brazil, and Mexico. Black points are occurrences.

**Table 2.** Evaluation metrics for prey distribution models used as biotic covariates in the harpy eagle distribution models. All models selected with ΔAICc = 0.0. FC = feature classes: Linear (L) and Quadratic (Q), RM = regularization multiplier. OR10 = 10% training presence omission rate threshold. CBI = Continuous Boyce Index, pROC = partial Receiver Operating Characteristic ratios.

| Food resource SDM | FC | RM | OR10 | CBI | pROC |
| --- | --- | --- | --- | --- | --- |
| Capuchin monkey | LQ | 1.0 | 0.104 | 0.840 | 1.344 |
| Howler monkey | LQ | 1.0 | 0.104 | 0.969 | 1.509 |
| Three-toed sloth | LQ | 1.0 | 0.136 | 0.900 | 1.621 |
| Tree porcupine | LQ | 1.0 | 0.111 | 0.906 | 1.489 |
| Two-toed sloth | LQ | 1.5 | 0.109 | 0.932 | 1.802 |

### Harpy eagle distribution models

All four best-fit harpy eagle models (ΔAIC_c_ = 0.0) had feature classes Linear and Quadratic and a regularization multiplier β = 1. Optimal selected models had robust discrimination ability with omission rates (OR10) at expected values (Table 3). The A+B model had the highest model calibration performance but all models had high calibration accuracy between predicted environmental suitability and test occurrence points (range: CBI = 0.842—0.899). All models were robust against random expectations (range: pROC = 1.346-1.460). Visually, including prey distributions in both the B and A+B models constrainted harpy eagle distribution (Fig. 2), compared to using solely abiotic covariates. The B and A+B models captured more detail in defining areas of highest suitability and relative abundance for the harpy eagle (Fig. 2). This was noticeable especially across key areas of the harpy eagle range in Guyana, eastern Colombia, Panama and northern Peru and the central Amazon basin in Brazil.

**Figure 2.**
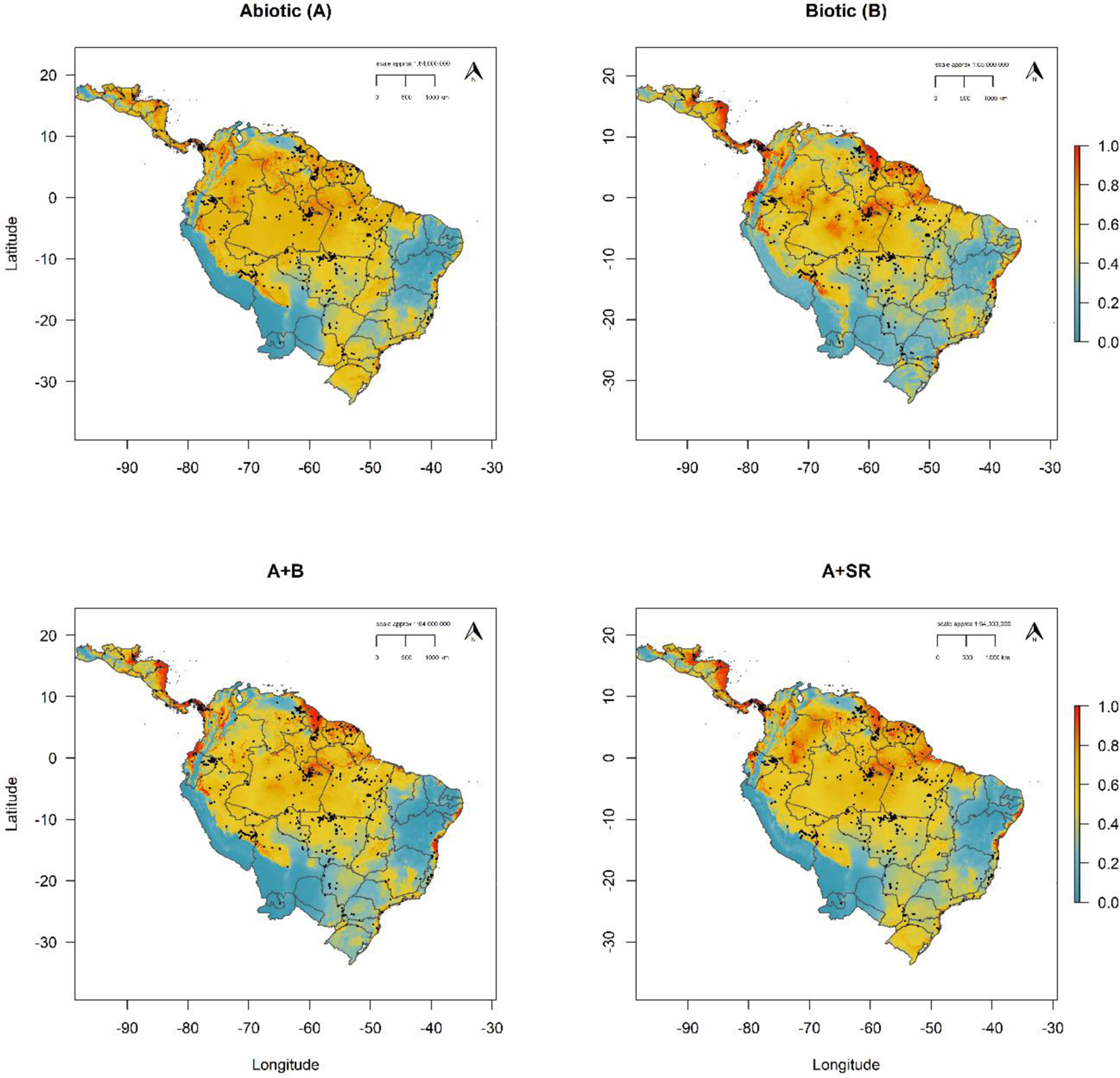
Predicted continuous distributions for the harpy eagle using abiotic and biotic covariates. Maps denote cloglog prediction with red areas (values closer to 1) having higher environmental suitability. Grey borders represent national borders and state boundaries for Argentina, Brazil, and Mexico. Black points define harpy eagle occurrences.

**Table 3.** Model selection and evaluation metrics for all four harpy eagle SDMs ranked by AICc. Evaluation metrics are Continuous Boyce Index (CBI) and tested against null expectations using partial Receiver Operating Characteristic ratios (pROC). OR10 = 10% training presence omission rate threshold.

| SDM | $AIC_c$ | OR10 | CBI | pROC |
| --- | --- | --- | --- | --- |
| A+B | 18027.82 | 0.102 | 0.899 | 1.460 |
| A+SR | 18095.91 | 0.098 | 0.842 | 1.414 |
| A | 18119.88 | 0.097 | 0.895 | 1.346 |
| B | 18177.30 | 0.101 | 0.872 | 1.399 |

### Predictor importance and responses

Climatic Moisture Index (CMI) contributed the highest percentage to the Abiotic model prediction (80.0 %), with three-toed sloth the highest contributor in both Biotic (71.3 %) and A+B (57.1 %) models. Species richness was the most important predictor (56.6 %) in the A+SR model, followed by CMI (35.2 %, Table S4). Response curves for covariates in the Abiotic model (Fig. 3) showed a unimodal response to CMI peaking at 0.4, with a positive response to minimum temperature for the warmest month peaking at suitable temperatures of 27°C. Harpy eagle occurrence had a consistently high positive response to higher predicted values of three-toed sloth occurrence in both the Biotic and A+B models (Figs. 4-5), with positive sigmoidal responses to species richness in both the Biotic and A+SR models (Fig. 6).

**Figure 3.**
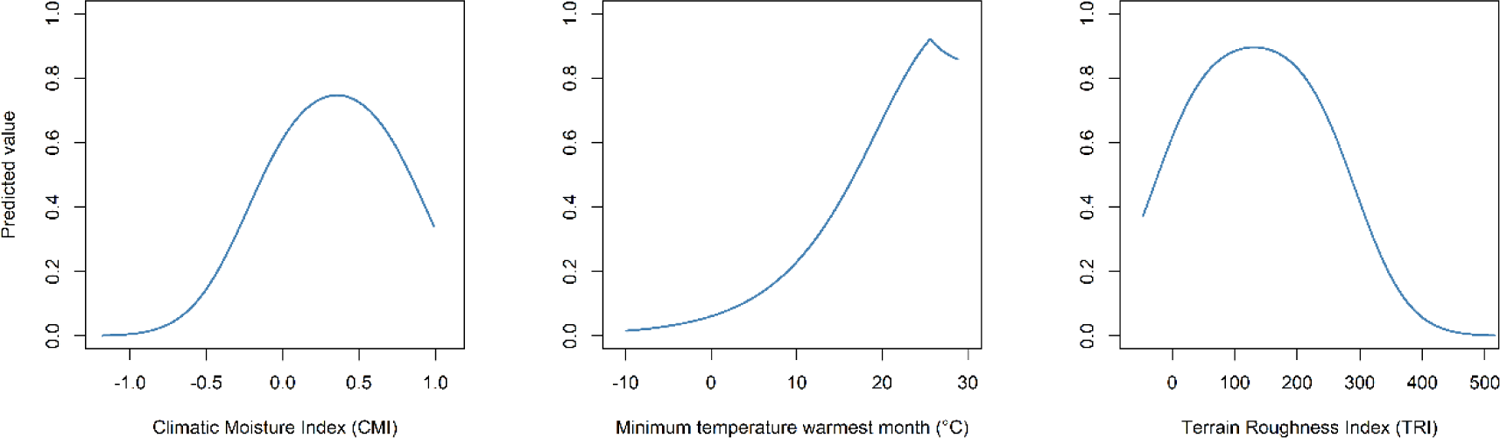
Response curves for covariates in the Abiotic distribution model for the harpy eagle. The response curves show the contribution to model prediction (y-axis) as a function of each continuous habitat covariate (x-axis). Maximum values in each response curve define the highest predicted relative suitability. The response curves reflect the partial dependence on predicted suitability for each covariate and the dependencies produced by interactions between the selected covariate and all other covariates.

**Figure 4.**
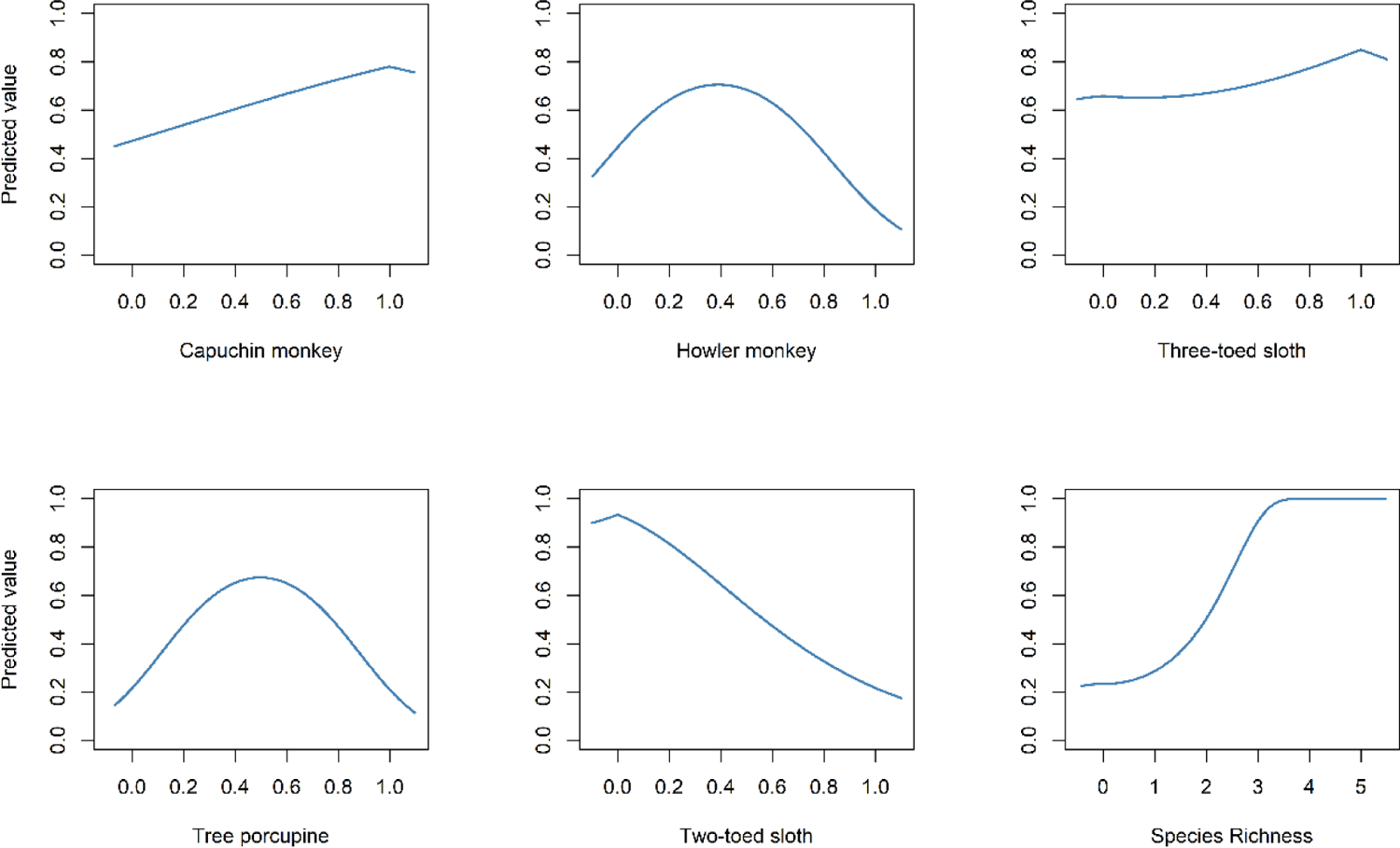
Response curves for covariates in the Biotic distribution model for the harpy eagle. The response curves show the contribution to model prediction (y-axis) as a function of each continuous habitat covariate (x-axis). Maximum values in each response curve define the highest predicted relative suitability. The response curves reflect the partial dependence on predicted suitability for each covariate and the dependencies produced by interactions between the selected covariate and all other covariates.

**Figure 5.**
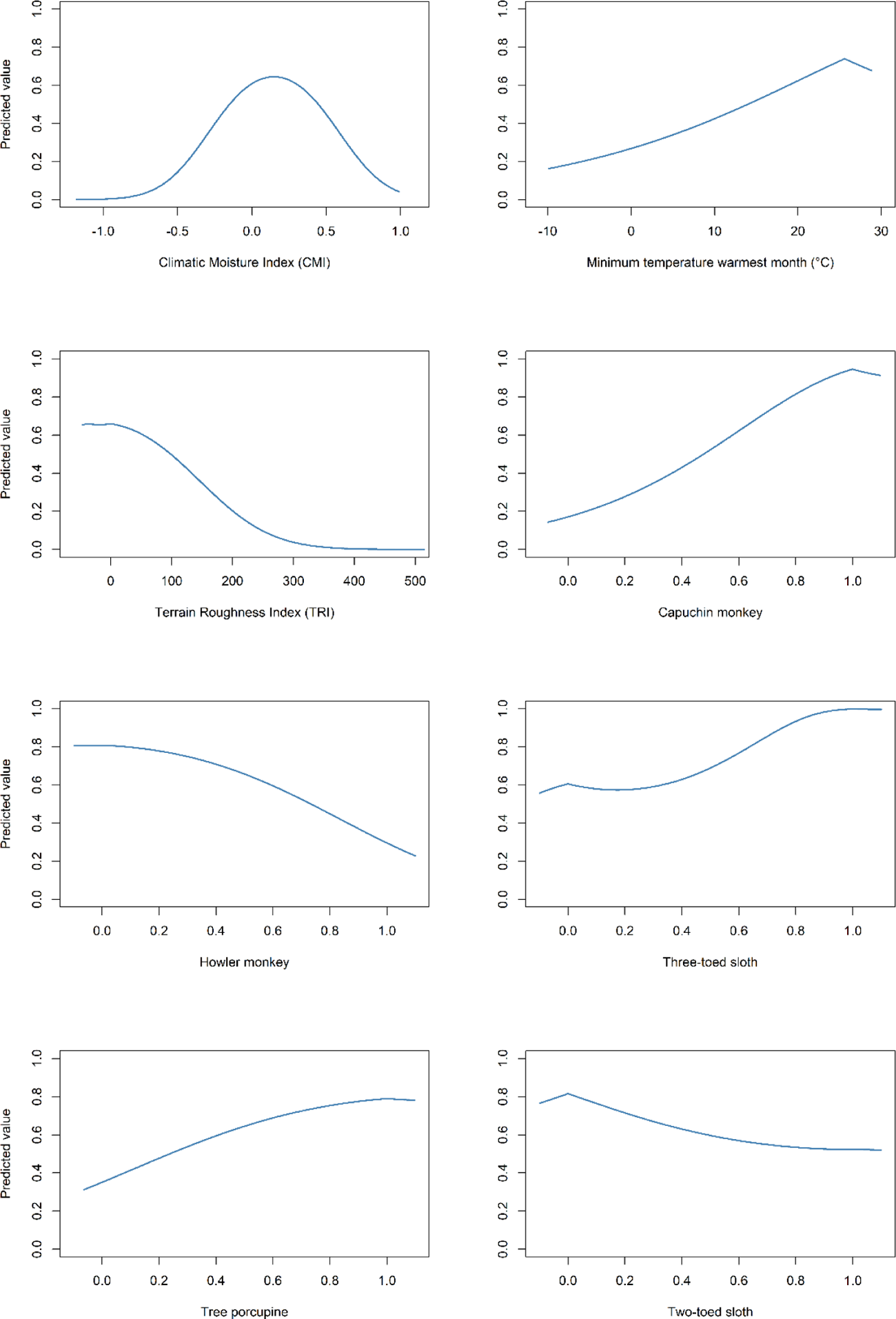
Response curves for covariates in the A+B distribution model for the harpy eagle. The response curves show the contribution to model prediction (y-axis) as a function of each continuous habitat covariate (x-axis). Maximum values in each response curve define the highest predicted relative suitability. The response curves reflect the partial dependence on predicted suitability for each covariate and the dependencies produced by interactions between the selected covariate and all other covariates.

**Figure 6.**
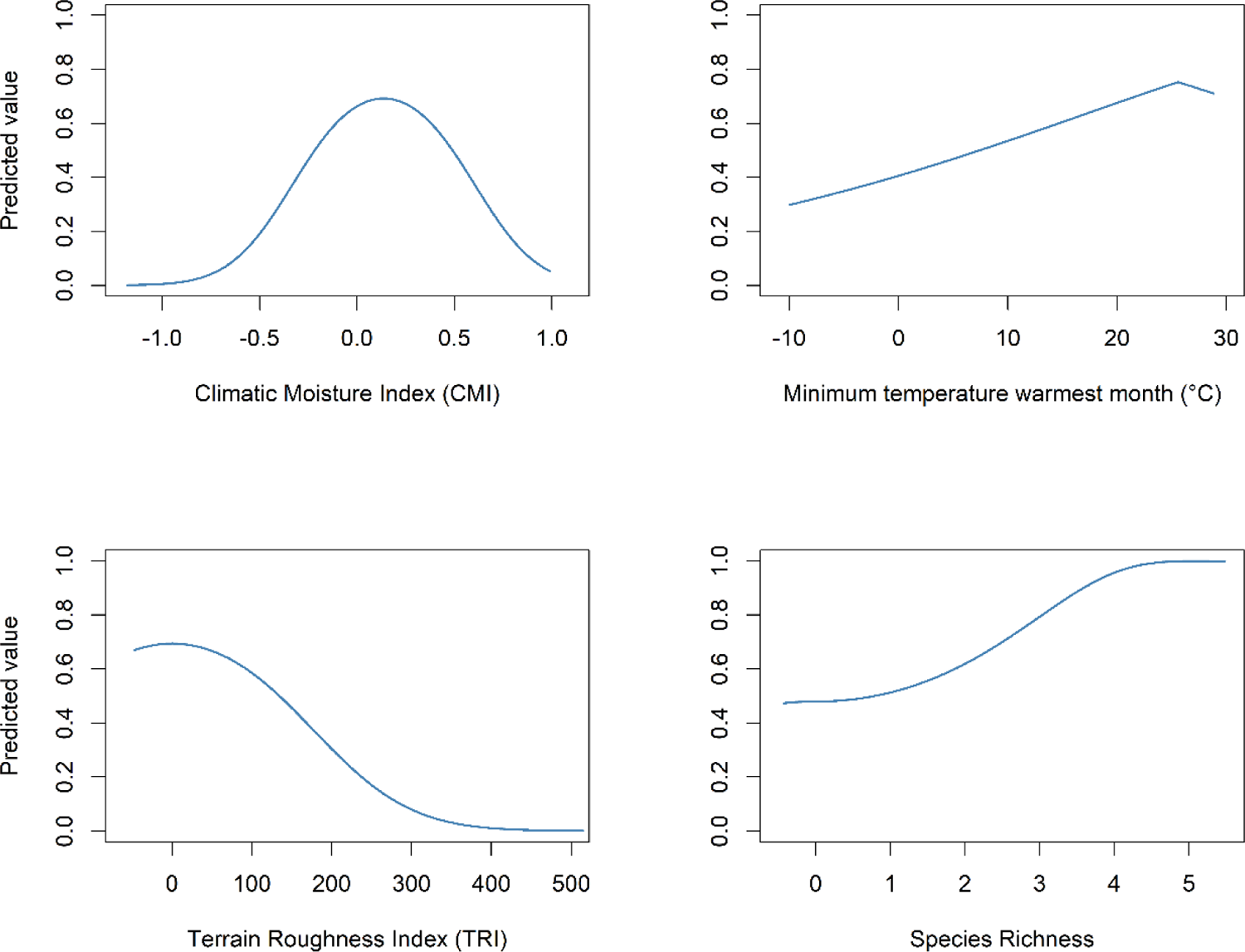
Response curves for covariates in the A+SR distribution model for the harpy eagle. The response curves show the contribution to model prediction (y-axis) as a function of each continuous habitat covariate (x-axis). Maximum values in each response curve define the highest predicted relative suitability. The response curves reflect the partial dependence on predicted suitability for each covariate and the dependencies produced by interactions between the selected covariate and all other covariates.

Harpy eagle occurrence responses to food resource distribution varied between models. There were positive responses to capuchin monkey occurrence in both the Biotic and A+B models, but with negative responses to two-toed sloth distribution, especially in the Biotic model (Figs. 4-5). Howler monkey and tree porcupine had unimodal responses in the Biotic model, peaking at ∼0.4 (Fig. 4). In the A+B model, these responses changed with a positive response to tree porcupine occurrence and a negative response to howler monkey occurrence (Fig. 5). The Abiotic predictor responses largely remained unchanged, when including biotic covariates, except for the sigmoidal response to terrain roughness (TRI) in both A+B models, compared to the unimodal response in the Abiotic model.

Model parameter estimates showed positive linear relationships with CMI in all three models using Abiotic covariates, but negative quadratic relationships (Table 4). Minimum temperature of the warmest month and TRI coefficients were shrunk to zero and showed no linear or quadratic relationships when biotic covariates were included. Both sloth genera had positive quadratic relationships in the Biotic model, with the relationship stronger in the A+B model, though two-toed sloth had a negative linear relationship. Tree porcupine had the strongest linear relationship in the Biotic model, followed by howler monkey, but these responses were less pronounced in the A+B model.

**Table 4.** Linear and Quadratic (with superscript 2) parameter estimates for each optimal model derived from penalized elastic net regression beta coefficients.

| Predictor | Abiotic | Biotic | A+B | A+SR |
| --- | --- | --- | --- | --- |
| Climatic Moisture Index | 1.96 |  | 1.37 | 0.99 |
| Min. temp. warmest month | 0.14 |  | 0.01 | 0.00 |
| Terrain Roughness Index | 0.02 |  |  | 0.00 |
| Climatic Moisture Index<sup>2</sup> | -3.35 |  | -4.70 | -3.89 |
| Min. temp. warmest month<sup>2</sup> |  |  |  | 0.00 |
| Terrain Roughness Index<sup>2</sup> | 0.00 |  | 0.00 | 0.00 |
| Tree porcupine |  | 6.47 | 1.75 |  |
| Two-toed sloth |  | -3.07 | -2.58 |  |
| Howler monkey |  | 2.60 | 0.00 |  |
| Capuchin monkey |  |  | 1.71 |  |
| Tree porcupine<sup>2</sup> |  | -6.90 |  |  |
| Howler monkey<sup>2</sup> |  | -4.03 | -1.38 |  |
| Capuchin monkey<sup>2</sup> |  | -0.93 | 0.69 |  |
| Two-toed sloth<sup>2</sup> |  | 0.66 | 1.47 |  |
| Three-toed sloth<sup>2</sup> |  | 0.52 | 2.13 |  |
| Species richness<sup>2</sup> |  | 0.27 |  | 0.10 |

### Geographic overlap and correlation

In geographic space, pair-wise overlaps between the harpy eagle and its food resource distributions were highest with capuchin (*D* = 0.867) and howler monkey (*D* = 0.858), followed by tree porcupine (*D* = 0.814). Three-toed sloth (*D* = 0.680) and two-toed sloth (*D* = 0.635) both had similar, but lower overlap scores compared to the primate and porcupine genera. The most correlated areas of distribution were first along the Caribbean coast of Central America, extending into the Chocó region along the Pacific coast of Colombia (Fig. 7). Second, a large but patchy area of high environmental suitability was predicted across Amazonia, extending from eastern Colombia, across the Guiana Shield and south into the northern Amazon of Brazil.

**Figure 7.**
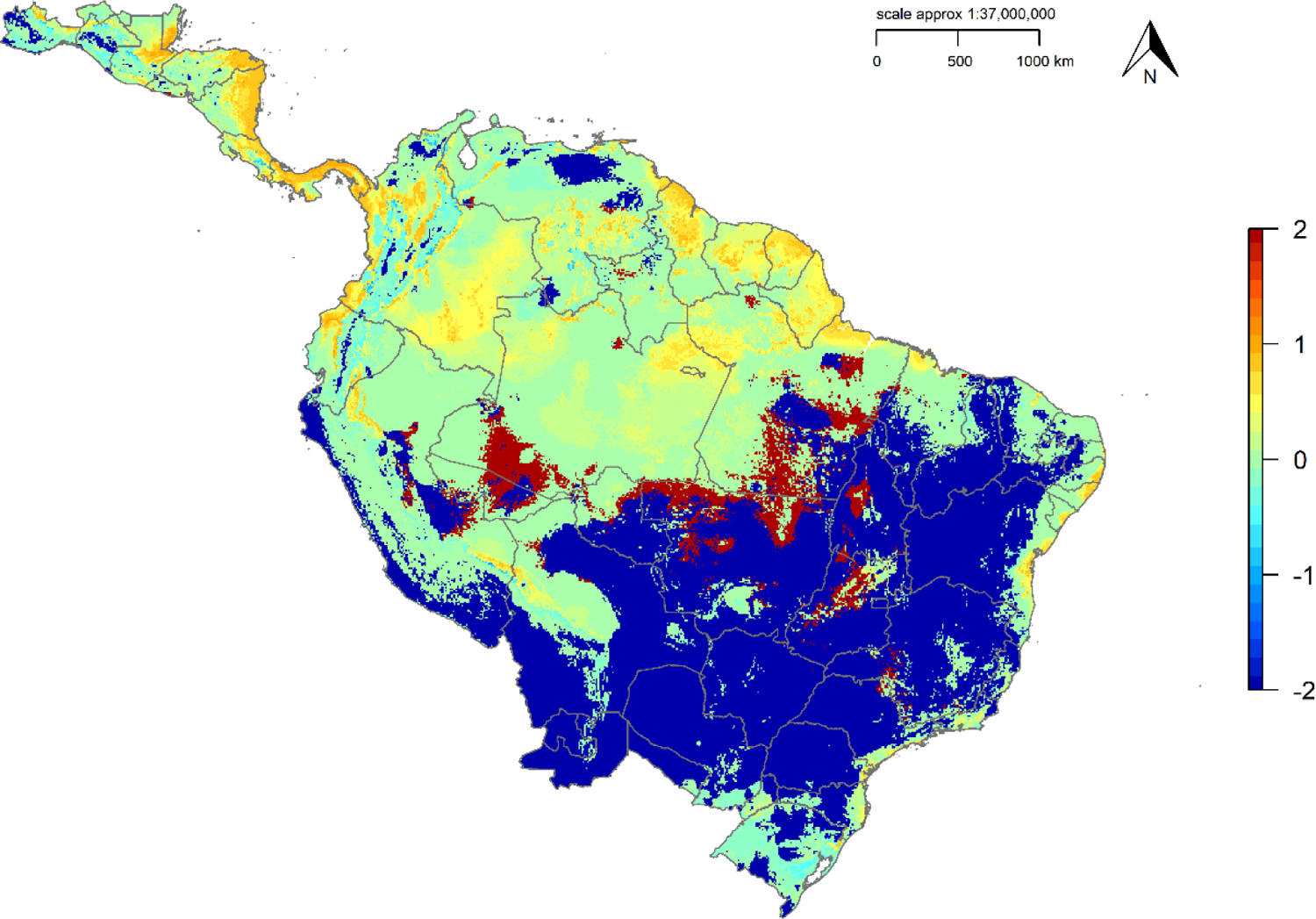
Predicted distribution correlation for the harpy eagle given the distribution of its five main prey species. Values close to −2 suggest absence, −1 to 0 can be interpreted as colonisable areas, 0 to 1 defines areas of highest suitability (prey availability) and values of 2 (dark red patches) show the most unsuitable (low prey availability) areas.

### Environmental overlap

In environmental space the core of prey distribution overlapped with the main core area for harpy eagle distribution, but the harpy eagle also occurs outside of this core food resource range (Fig. 8). Measuring overlap in environmental space resulted in moderate overlap (*D* = 0.424), with the hypothesis of equivalence between the harpy eagle and its food resource distribution rejected from the Equivalency (*P* = 0.02) and Background Statistics (*P* = 0.27). Both the harpy eagle and its main prey occupied more dissimilar environmental space than expected by chance, with the food resource distribution occupying a more restricted environmental space compared to the harpy eagle (Figs. S1-S2).

**Figure 8.**
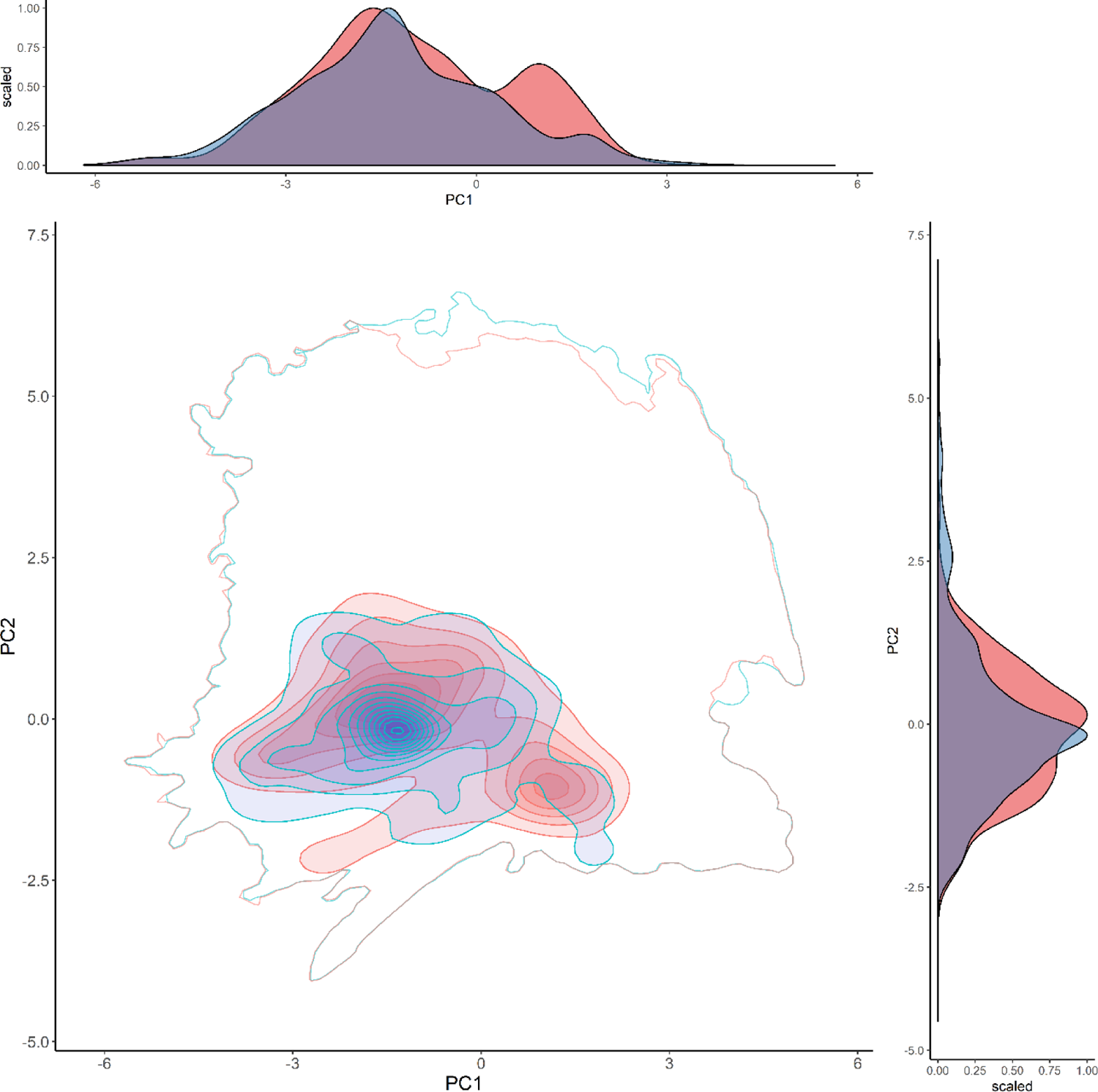
Environmental overlap (purple) for the harpy eagle (red) and its five main prey species combined (blue) across environmental space from the first two principal components.Total variance explained by the two principal components = 62.71 % (PC1 = 41.72 %, PC2 = 20.99 %). Filled isopleths are kernel densities from 1-100%. Empty kernel density isopleths represent 1% density isopleth of the environment.

## Discussion

Recent theoretical and empirical work has demonstrated the importance of including resource distributions in macro-scale SDMs (Araújo & Rozenfeld 2014; Atauchi *et al*. 2018; Ghergel *et al*. 2018; Palacio & Girini 2018). Our results show that incorporating the distribution of the harpy eagle’s five main prey species at a continental scale improved its distribution estimates compared to using solely abiotic covariates. This result further counters the Eltonian Noise Hypothesis (Soberón & Nakamura 2009), the assumption that biotic interactions are unimportant at broad spatial scales (Pearson & Dawson 2003). Including food resources as individual prey species distribution rasters improved the predictive performance of the Abiotic model. Moreover, using solely biotic covariates or combined as species richness still resulted in high performing models, but the combination of A+B was best. Geographic overlap ranged from moderate to high between the harpy eagle and its main prey species. However, overlap was lower in environmental space when all main prey occurrences were combined, suggesting the harpy eagle switches to other food sources outside of core areas that enable existence in peripheral habitats.

The spatial pattern of species’ distributions are products of physiological constraints such as climate and topography, and interactions with other co-occurring species (MacArthur 1972). It follows then that both abiotic and biotic factors combined should drive species distributions, and abiotic variables alone are unable to provide sufficient detail for distribution estimates at coarse scales (Wisz *et al*. 2013; Kass *et al*. 2019). Our results support this conclusion by improving an abiotic model prediction with the inclusion of food resource distributions. Three-toed sloth was the most important biotic predictor in both the Biotic and A+B models (Table S4), consistent with this species being the principal prey for the harpy eagle across its range (Aguiar-Silva *et al*. 2014; Miranda 2015; Miranda 2018). However, the importance of three-toed sloth distribution decreased when including abiotic factors, with CMI the second most important predictor in the A+B model. This indicates that only a reduced subset of climatic and biotic covariates are necessary to account for the major distributional constraints for the harpy eagle.

In the A+SR model, species richness was the most important predictor (56.6 %), followed by CMI (35.2 %). Combined, these two covariates accounted for nearly 92 % of model prediction, further supporting the inclusion of food resource species richness as a predictor in SDMs. Yet, when including species richness in the Biotic model its importance was low, probably due to any predictive power lost amongst the other biotic noise from the individual food resource covariates. For SDMs food resource distributions should thus be included as single covariates where predator-prey interactions are well established, and occurrence data are available. However, if occurrence data for single prey species are lacking (as is often the case), then combining all known food resource species into a single predictor is a valid method (Kass *et al*. 2019). One possible limitation of this study was only including the five main prey species when it is known that other prey species are consumed by the harpy eagle across its range (Miranda 2015). Indeed, the occurrence of suitable environmental space for the harpy eagle outside of its main food resource distributions suggests switching to other food types outside of its core range.

Our results confirm the importance of sloth distribution as one of the main drivers for harpy eagle distribution. There were high positive responses between harpy eagle distribution and three-toed sloth occurrence and both sloth genera had the highest percent contributions to the Biotic model prediction. Indeed, in some parts of their range harpy eagles have narrow diets comprised of 80 to 95 % sloths (Miranda *et al*. 2020), in central and eastern Amazonia (Galetti & de Carvalho 2000; Aguiar-Silva *et al*. 2014) and north-east Ecuador (Muñiz-López 2008). However, the harpy eagle is not so specialized on a diet of sloths as to be absent from areas where sloths are not present. It seems likely that in the southern and eastern parts of the harpy eagle range primates and porcupines are the key prey species, replacing sloths as the primary food source (Miranda 2015). Thus, our models are able to capture the spatial variation in predator-prey distribution across a continental tropical forest system by using a range of key prey genera and not relying solely on a single biotic predictor.

Using response curves to interpret model outputs is a useful though underused aspect of model evaluation in many SDMs (Guevara *et al*. 2018; Kass *et al*. 2019). Here, modelled responses for the three-toed sloth were strongly positive in both the Biotic and A+B models, peaking at 1.0 as expected (Figs. 4-5). Capuchin monkey followed similar positive responses, yet there was a flatter negative response to two-toed sloth distribution. Given the broad distribution of the two-toed sloth closely matching that of the harpy eagle this seems counter-intuitive, and it is difficult to explain this response. Both howler monkey and tree porcupine had unimodal responses in the Biotic model, peaking at 0.4. However, when all covariates were included in the A+B model, tree porcupine showed a clear positive response. Conversely, harpy eagle distribution then showed a negative response to howler monkey distribution, opposite to the positive response in the Biotic model. This variation in model responses suggests either prey switching or highly complex interactions occurring between abiotic and biotic processes. Using methods that are capable of modelling complex species interactions (e.g. regression-based, Aragón & Sánchez-Fernández 2013; or Bayesian networks, Staniczenko *et al*. 2017) may be a way forward to tackle this problem for biotic SDMs.

Pair-wise geographic overlaps supported strong relationships in distribution between the harpy eagle and its main food resources. High overlaps with most of its main prey suggests harpy eagle distribution is largely dependent on where its main food resources exist. Both primate prey genera (capuchin and howler monkey) had higher overlap values than the other main prey species. This could be partly explained by both primate genera having similar broad distributions across the Neotropics to the harpy eagle, thus overlap values would be expected to be high. Conversely, overlaps for both sloth genera were lower, even though in many areas of the harpy eagle range sloths are often the primary food resource. However, both sloth genera have more restricted ranges than both the primate genera, thus overlap values would be expected to be lower. The correlation model predicted the most common areas of distribution across Amazonia, the Guiana Shield, and the Caribbean coast of Central America. Given the high reliance that harpy eagle distribution has with its main food resources, we recommend conserving tropical lowland forest habitat and prioritizing research in these regions.

Interestingly, when all food resources were combined in environmental space, overlap was markedly lower than any single overlap in geographical space. The harpy eagle and combined prey distributions were significantly different from each other in environmental space, suggesting that the harpy eagle is able to exist in areas outside of its main food resource distributions. Harpy eagles are known to feed on over 100 prey species and are adaptable in their food choices when required (Miranda 2015, 2018). Thus, even in areas where its main prey is not abundant, the harpy eagle is presumably still able to switch to other food types, expanding its potential distribution beyond its core range. Measuring distribution in environmental space is thus useful as a comparison to geographical space, giving a more in-depth picture across different dimensions in distribution (Sutton *et al*. 2022b). Complex distributional relationships between taxa can often go undetected in geographical space (Warren *et al*. 2019). Thus, comparing distributions of co-occurring taxa in environmental space provides a more comprehensive account of niche dimensions.

The biotic interactions hypothesis states that species interactions are the main driver for species distribution in the relatively stable climates of the tropics (MacArthur 1972; Louthan *et al*. 2015). Our results in general support this, though abiotic processes are clearly important, with Climatic Moisture Index (CMI) still the key abiotic predictor in the A+B and A+SR models. Because CMI is closely correlated with the primary vegetation types in Neotropical forests (Beck *et al*. 2018), it seems likely that CMI is acting as a proxy for lowland tropical forest, which by definition is the key vegetation type for all species distributions in this tropical forest system. Thus, both specific food resources and habitat type are likely the main drivers on harpy eagle distribution, which hardly seems unexpected. A useful next step would be to include direct habitat variables, competitor distributions and human impacts, along with food resources, to provide a broader perspective on the main influences determining harpy eagle distribution (Joint-SDMs, Pollock *et al*. 2014).

Consistent with previous smaller scale regional studies (e.g. Hof *et al*. 2012; Aragón *et al*. 2018; Ghergel *et al*. 2018), our results support including food resources in SDMs, but also that including the main food resource distributions for apex predators is important at continental scales. Further, our results dispute the Eltonian Noise Hypothesis, similar to conclusions from landscape to regional scale studies (Araújo *et al*. 2014; Atauchi *et al*. 2018). However, it is recognised that increases in predictive power were relatively slight and the Abiotic SDM still had high predictive accuracy. Including resource distributions has much practical value for advancing SDM predictions across a range of applications in space (spread of invasive species) and time (climate change range shifts). As demonstrated here, predictions were improved when applied to basic model interpolation, thus not including resource distributions may result in poorer model transferability when extrapolating in space and time.. However, we recognise limitations with interpreting model outputs, with the prey model for two-toed sloth over-predicting in northern Central America, beyond the species’ known northern range limits (McCarthy *et al*. 1999), which will have subsequent impacts on harpy eagle range predictions.

We show how incorporating food resource distributions improves model predictive power and circumscribes the spatial complexity in harpy eagle distribution. Adding food resource distributions revealed the crucial role of predator-prey interactions in harpy eagle distribution. Given the wide variation in food type taken by the harpy eagle across its range (Aguiar-Silva *et al*. 2014; Miranda 2018), maintaining these prey resources should also be a priority in conservation programs for the harpy eagle. Conserving habitat for the key arboreal mammal prey populations along with one of their main predators as a complete tropical forest system seems a viable approach given the reliance on harpy eagle presence with their main food resource distributions. We encourage practitioners to incorporate known biotic interactions into SDMs, but modellers should recognise that understanding the complex interactions inherent in natural systems is a challenge (Aragón *et al* 2018). Whilst we demonstrate that using resource distributions improves model predictions at macro-scales, this needs further testing across multiple taxa and ecosystems to determine if this finding is consistent elsewhere.

## Acknowledgements

We thank all individuals and organisations who contributed occurrence data to the Global Raptor Impact Network (GRIN) information system. LJS thanks The Peregrine Fund for providing financial assistance for his studentship. EBPM work is supported by Rufford Foundation, ONF Brasil, Rainforest Biodiversity Group, Idea Wild, Explorer’s Club Exploration Fund, Cleveland Metroparks Zoo, and SouthWild.com. We thank the M.J. Murdock Charitable Trust for funding and the Information Technology staff and Research Library interns at The Peregrine Fund for support.

## Authors’ contributions

L.J.S., M.F., C.J.W.M, & R.P. conceived the ideas and designed methodology; D.L.A, E.B.P.M., F.H.V. & J.J.V.G collected data; L.J.S. analysed the data; L.J.S. led the writing of the manuscript. All authors contributed critically to the drafts and gave final approval for publication.

## Data Availability Statement

Upon acceptance the data that support the findings of this study will be made openly available on the data repository *figshare*.

## Supplementary Information

## Appendix 1 Supplementary Tables

**Table S1.** Environmental variables used as covariates for food resource distribution models used as biotic covariates in the harpy eagle distribution models. Black points indicate which environmental variables were used in each respective species distribution model.

| Predictor | Capuchin monkey | Howler monkey | Three-toed sloth | Tree porcupine | Two-toed sloth |
| --- | --- | --- | --- | --- | --- |
| Mean diurnal temperature range |  |  | • |  | • |
| Isothermality | • | • | • |  |  |
| Mean temperature wettest quarter | • |  |  |  |  |
| Precipitation wettest month |  |  |  | • |  |
| Precipitation driest month | • | • |  |  |  |
| Precipitation warmest quarter | • | • | • | • | • |
| Precipitation coldest quarter |  |  |  |  |  |
| Climatic Moisture Index |  | • | • | • | • |
| Minimum temperature warmest month |  | • |  |  |  |
| Maximum temperature coldest month |  |  | • |  |  |
| PET driest quarter |  |  |  | • | • |
| PET seasonality |  |  |  |  |  |
| PET warmest quarter | • | • |  |  |  |
| PET wettest quarter | • | • | • | • | • |
| Topographic wetness |  |  |  |  |  |
| Terrain Roughness Index | • | • | • | • | • |

**Table S2.** Multi-collinearity test using stepwise elimination Variance Inflation Factor (VIF) for correlation between food resource distribution models used as biotic covariates.

| Resource distribution model | VIF |
| --- | --- |
| Two-toed sloth *Choloepus* | 9.260 |
| Three-toed sloth *Bradypus* | 8.732 |
| Capuchin monkey *Cebus* & *Sapajus* | 4.984 |
| Howler monkey *Alouatta* | 3.722 |
| Tree porcupine *Coendou* | 2.269 |

**Table S3.** Selection of variables for environmental overlap analysis using stepwise elimination Variance Inflation Factor (VIF) to reduce multi-collinearity between variables.

| Environmental variable | VIF |
| --- | --- |
| Climatic Moisture Index | 6.486 |
| Precipitation wettest month | 4.432 |
| PET wettest quarter | 3.724 |
| PET warmest quarter | 3.637 |
| Precipitation driest month | 2.933 |
| Precipitation warmest quarter | 2.684 |
| Mean diurnal temperature range | 2.037 |
| Isothermality | 1.899 |
| Terrain Ruggedness Index | 1.516 |

**Table S4.** Percent contribution to model prediction for environmental covariates used in all SDMs for the harpy eagle. Ranked by highest % contribution to the Abiotic and Biotic models.

| Predictor | Abiotic | Biotic | A+B | A+SR |
| --- | --- | --- | --- | --- |
| Climatic Moisture Index | 80.0 |  | 28.6 | 35.2 |
| Minimum temperature warmest month | 14.7 |  | 0.1 | 4.3 |
| Terrain Roughness Index | 5.3 |  | 5.1 | 3.9 |
| Three-toed sloth |  | 71.3 | 57.1 |  |
| Tree porcupine |  | 9.8 | 1.9 |  |
| Howler monkey |  | 8.6 | 1.8 |  |
| Capuchin monkey |  | 6.9 | 3.8 |  |
| Two-toed sloth |  | 2.6 | 1.5 |  |
| Species Richness |  | 0.8 |  | 56.6 |

## Appendix 2 Supplementary Figures

**Figure S1.**
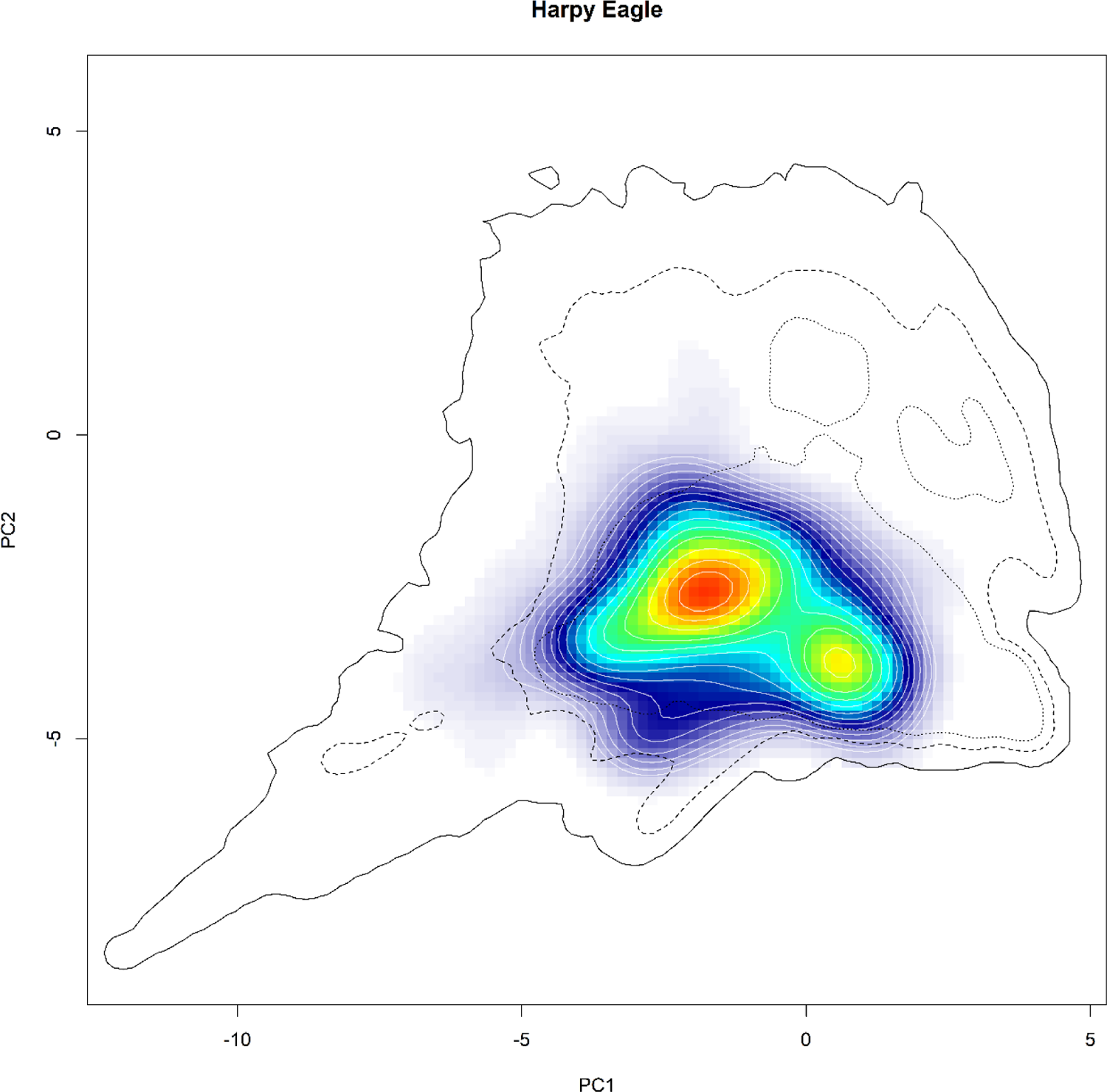
Distribution in environmental space for the harpy eagle across the first two principal components. Red areas indicate highest environmental suitability. Filled kernel density isopleths characterize kernel density values from 0.4 (blue) to 0.99 (red). Black isopleth lines define kernel density of the corresponding environment, with black solid line = 0.1, black hashed line = 0.5, black dotted line = 0.75.

**Figure S2.**
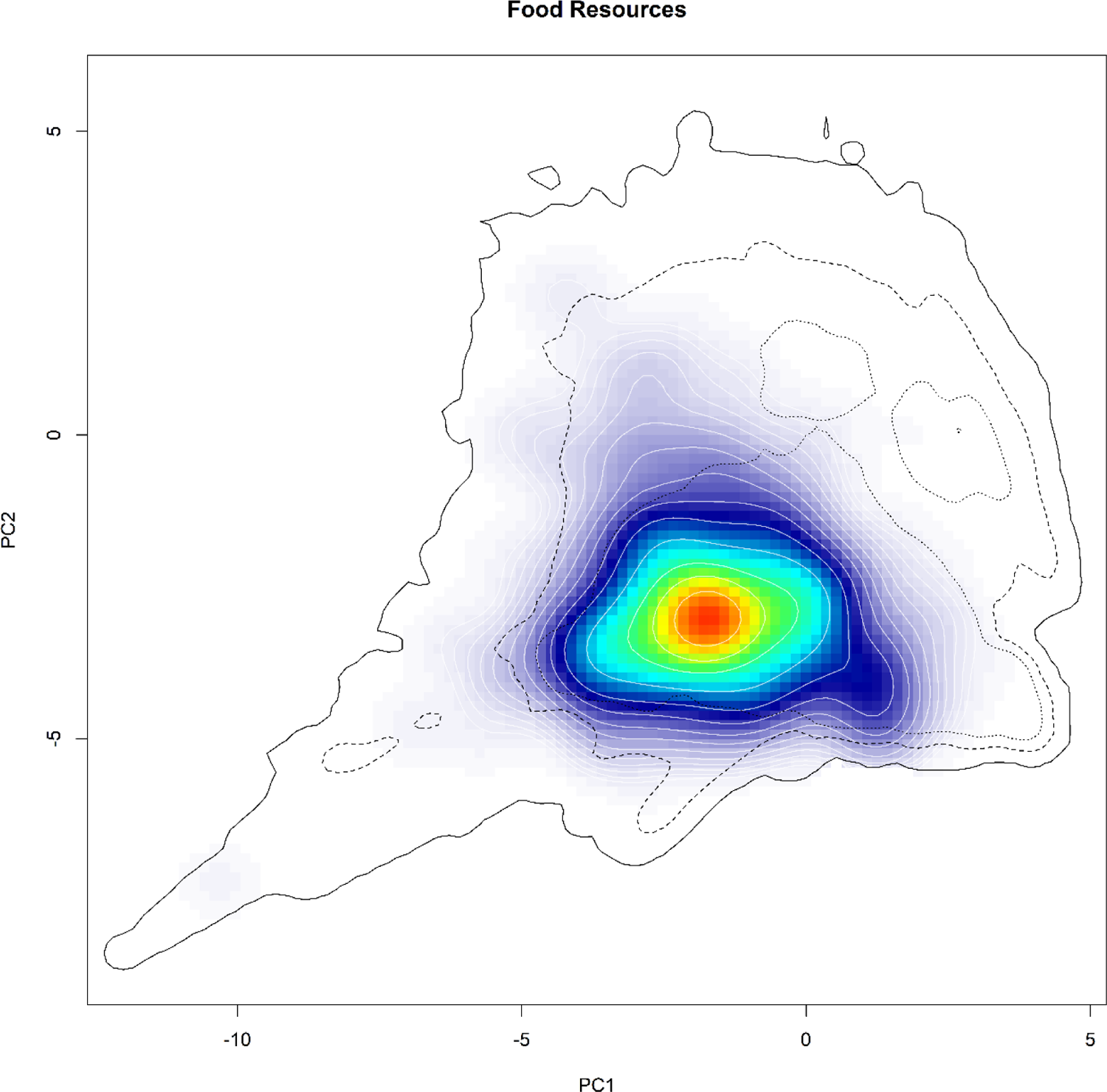
Distribution in environmental space for the food resource genera across the first two principal components. Red areas indicate highest environmental suitability. Filled kernel density isopleths characterize kernel density values from 0.4 (blue) to 0.99 (red). Black isopleth lines define kernel density of the corresponding environment, with black solid line = 0.1, black hashed line = 0.5, black dotted line = 0.75.

